# Dysfunctional T Follicular Helper Cells Cause Intestinal and Hepatic Inflammation in NASH

**DOI:** 10.1101/2023.06.07.544061

**Authors:** Haiguang Wang, Fanta Barrow, Gavin Fredrickson, Kira Florczak, Huy Nguyen, Preethy Parthiban, Adam Herman, Oyedele Adeyi, Christopher Staley, Sayeed Ikramuddin, Hai-Bin Ruan, Stephen C. Jameson, Xavier S. Revelo

## Abstract

Nonalcoholic steatohepatitis (NASH), characterized by hepatic inflammation and cellular damage, is the most severe form of nonalcoholic fatty liver disease and the fastest-growing indication for a liver transplant. The intestinal immune system is a central modulator of local and systemic inflammation. In particular, Peyer’s patches (PPs) contain T follicular helper (Tfh) cells that support germinal center (GC) responses required for the generation of high-affinity intestinal IgA and the maintenance of intestinal homeostasis. However, our understanding of the mechanisms regulating mucosal immunity during the pathogenesis of NASH is incomplete. Here, using a preclinical mouse model that resembles the key features of human disease, we discovered an essential role for Tfh cells in the pathogenesis of NASH. We have found that mice fed a high-fat high-carbohydrate (HFHC) diet have an inflamed intestinal microenvironment, characterized by enlarged PPs with an expansion of Tfh cells. Surprisingly, the Tfh cells in the PPs of NASH mice showed evidence of dysfunction, along with defective GC responses and reduced IgA^+^ B cells. Tfh-deficient mice fed the HFHC diet showed compromised intestinal permeability, increased hepatic inflammation, and aggravated NASH, suggesting a fundamental role for Tfh cells in maintaining gut-liver homeostasis. Mechanistically, HFHC diet feeding leads to an aberrant increase in the expression of the transcription factor KLF2 in Tfh cells which inhibits its function. Thus, transgenic mice with reduced KLF2 expression in CD4 T cells displayed improved Tfh cell function and ameliorated NASH, including hepatic steatosis, inflammation, and fibrosis after HFHC feeding. Overall, these findings highlight Tfh cells as key intestinal immune cells involved in the regulation of inflammation in the gut-liver axis during NASH.

## INTRODUCTION

Non-alcoholic fatty liver disease (NAFLD) is the most common form of liver disease worldwide, currently affecting more than 25% of Americans^1, 2^. As obesity and type 2 diabetes rates continue to surge, the prevalence of NAFLD has also increased, leading to an alarming health concern^3^. NAFLD is a progressive condition that initiates with hepatic steatosis but that can evolve into a more severe disease entity known as non-alcoholic steatohepatitis (NASH), which presents features of hepatocellular injury and fibrosis^2, 4^. It is estimated that nearly 25% of adults with NAFLD have NASH^2, 4^ with an increased risk of developing debilitating conditions such as cirrhosis and hepatocellular carcinoma^2, 4^. Despite recent efforts, our understanding of the precise cellular and immune mechanisms orchestrating NASH pathogenesis remains incomplete.

Human studies have implicated changes in the gut microbiota in promoting NAFLD and NASH^5–10^, through mechanisms including increased gut permeability and intestinal bacterial overgrowth^11^. Disruptions in the commensal microbiome can lead to altered metabolite production, such as decreased levels of butyrate, critical for maintaining intestinal integrity permeability^12, 13^. A compromised intestinal integrity, in turn, allows the translocation of endotoxin, microbial antigens, and other inflammatory factors from the gut into the liver, leading to hepatic inflammation^6–9^. Previous studies have demonstrated that the adaptive immune system is critical in regulating the microbiota and intestinal homeostasis, particularly through the actions of immunoglobulin A (IgA)^14–16^. IgA is the most abundantly produced antibody in mammals^17, 18^ that directly targets gut microbes to restrain the outgrowth of specific microbes and promote their diversity^17^. Consequently, a reduction in the affinity or specificity of intestinal IgA towards gut microbes can compromise this balance^16^, and may influence NASH.

Peyer’s patches (PPs) are part of the gut-associated lymphoid tissue (GALT) distributed throughout the small intestine^19, 20^ and responsible for the regulation of the intestinal immune responses^19, 20^. Indeed, PPs act as a central hub for the initiation of the adaptive immune response in the gut^19, 20^. The PPs harbor germinal center (GC) responses, critical for the affinity maturation of antibodies, the generation of memory B cells and long-lived plasma cells, and the production of intestinal IgA^19–21^. In the GC, the interaction between Tfh cells and B cells is essential for somatic hypermutation and class switching recombination ^22–24^. However, the role and mechanisms by which Tfh cells regulate obesity-related metabolic diseases, such as NASH, remains inadequately understood.

Here, we focused on the role of intestinal Tfh cells in the maintenance of gut homeostasis and disease progression during HFHC diet-induced NASH. We found that intestinal Tfh cells expanded in the PPs of NASH mice where they play an essential role in maintaining gut homeostasis and mitigating liver inflammation. Furthermore, intestinal Tfh cells exhibit functional impairments during NASH, leading to diminished GC responses in the intestine and a subsequent decrease in gut IgA^+^ B cell responses. Mechanistically, NASH results in an aberrant upregulation of Kruppel-like factor 2 (KLF2) in Tfh cells while its insufficiency reverses the dysfunctional phenotype of Tfh cells and mitigates NASH progression. Overall, our findings shed light on the fundamental role of intestinal Tfh cells in maintaining gut homeostasis and highlight the pivotal role played by intestinal immunity in the pathogenesis of NASH.

## RESULTS

### Intestinal Tfh cells expand in the PPs during NASH

We have previously reported that HFHC feeding for 20 weeks leads to severe NASH including obesity hepatic steatosis, inflammation, and fibrosis^5^ (Supplementary Figure 1A). To determine how NASH influences the immune cell populations in intestinal PPs, we fed C57BL6/J mice either a normal chow diet (NCD) or a HFHC diet for up to 20 weeks. Histological analysis revealed enlarged PPs with increased cellularity in HFHC intestinal sections, compared with NCD controls (Figure 1A-B). Quantification of total CD45^+^ cells in PPs by flow cytometry confirmed the increased leukocyte number in HFHC mice (Figure 1C). While no differences were detected in the lamina propria (LPL), we also observed a substantial increase in cellularity in the mesenteric lymph node (mLN) and spleen of HFHC mice (Supplementary Figure 1B-D). Consistent with previous reports ^25, 26^, however, we detected increased Th17 cells in the LPL and mLN, but not in the spleen (Supplementary Figure 1E-G). To investigate the transcriptional landscape of PPs during NASH, we performed bulk RNA sequencing (RNA-seq) of PPs and mLNs isolated from NCD and HFHC mice. Principal component analysis (PCA) revealed a substantial separation between PPs from HFHC and NCD mice, while mLNs from both groups clustered together, indicating that NASH resulted in an altered gene expression profile unique to the PPs (Figure 1D). Gene set enrichment analysis demonstrated that PPs from HFHC mice were enriched in immune transcripts, particularly in B and T cell signatures (Figure 1E). To investigate the nature of such immune response, we determined the expression of a curated list of adaptive immune cell genes and found an increased expression of B cell and T cell genes in HFHC PPs (Figure 1F). Notably, PPs from HFHC mice showed an increased expression of Tfh cell-associated genes (*Bcl6*, *Cxcr5*, *Tox2*, *Icos*), but no difference in CD8 or other CD4 T cell subsets (Figure 1F). We confirmed the increase in B cells and CD4 T cells, but no changes in CD8 T cells in the PPs from HFHC mice by flow cytometry (Supplementary Figure 1H-J). Considering the Tfh gene signature observed in the PPs of HFHC mice, we further characterized the Tfh populations by flow cytometry, using an established gating strategy (Supplementary Figure 1K). We found that CD4 T cells from HFHC PPs show an elevated expression of CXCR5 (Figures 1G-I) and contain an increased frequency and cell number of Tfh and Pre-Tfh cells (Figure 1J-L). Collectively, these data demonstrate that NASH induces an increased immune response in PPs, characterized by an altered transcriptional landscape and a bias towards increased Tfh cell differentiation, suggesting a role for these cells in modulating intestinal immune responses during NASH.

**Figure 1.**
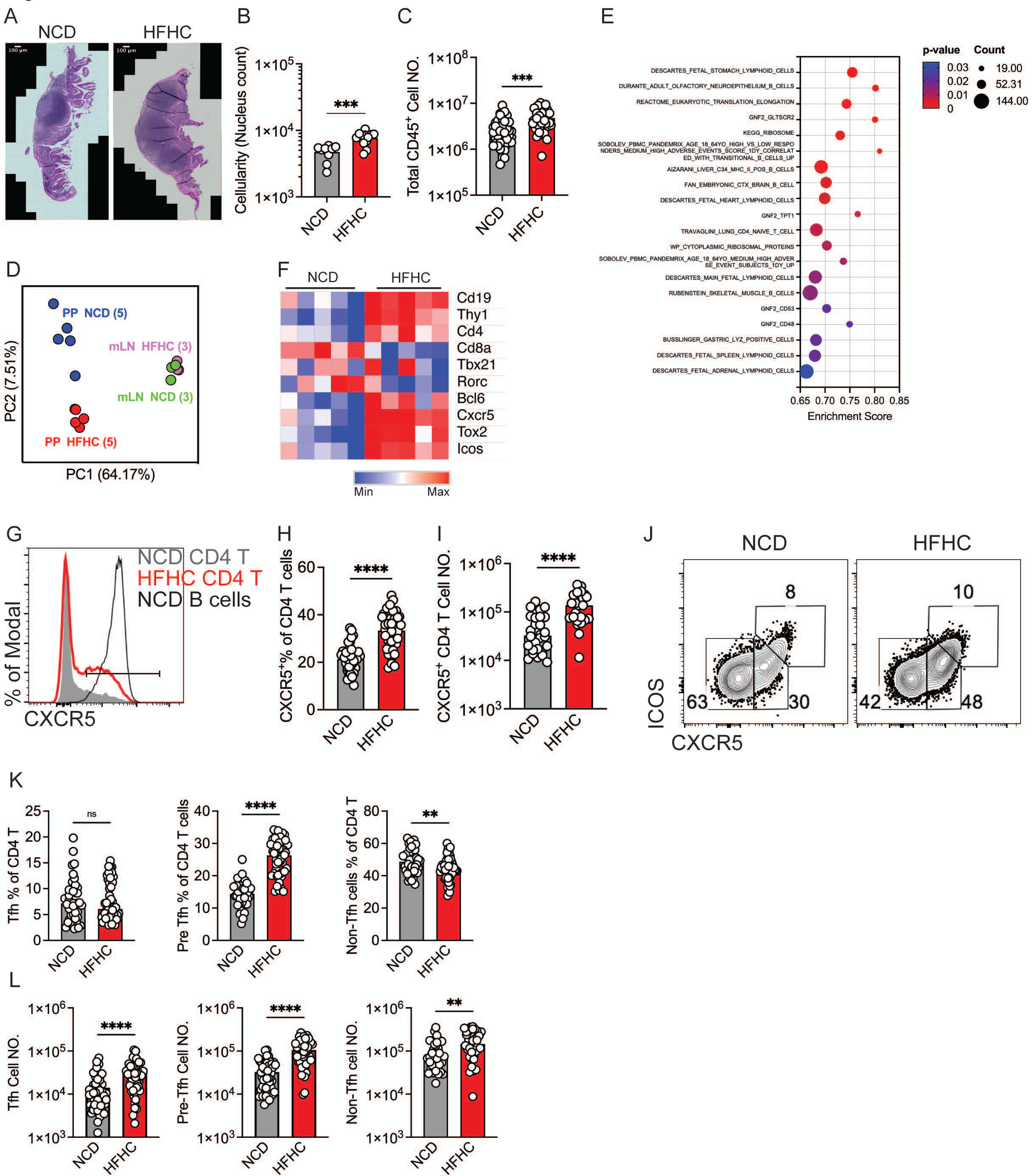
Enlarged PPs and expanded intestinal Tfh cells in HFHC diet-induced NASH mice. (A) Histological H&E staining of PPs from NCD and HFHC mice. (B) Cellularity (nucleus count) of PPs in H&E staining. (C) Cell number of total live CD45+ cells in PPs by flow cytometry. (D) Principal component analysis (PCA) of bulk RNA-seq data of PPs and mLN from NCD and HFHC mice. (E) Gene set enrichment analysis (GSEA) of bulk RNA-seq data of PPs (HFHC vs NCD), with Hallmark gene sets. (F) Heatmap of curated list of genes in PPs. (G) Flow cytometry histogram of CXCR5 expression in NCD CD4 T cell, HFHC CD4 T cell, NCD B cells of PPs. (H) Frequency of CXCR5^+^ CD4 T cells in PPs of NCD and HFHC mice. (I) Cell number of CXCR5^+^ CD4 T cells in PPs of NCD and HFHC mice. (J) Representative flow cytometry analysis to identify Tfh (CXCR5^high^ ICOS^high^), Pre-Tfh (CXCR5^intermediate^ ICOS^intermediate^) and Non Tfh (CXCR5^−^ ICOS^−^) cells among Foxp3^−^ CD4 T cells, in PPs of NCD and HFHC mice. (K) Frequency of Tfh (Left), Pre-Tfh (Middle) and Non-Tfh cells (Right) among total CD4 T cells. (L) Cell number of Tfh (Left), Pre-Tfh (Middle) and Non-Tfh cells (Right). Data are representative of three independent experiment, ns=not significant, ****p<0.001, **p<0.01, unpaired t test.

### Diminished Germinal Center and IgA Responses in NASH Mice

To further examine the effect of NASH on intestinal immune homeostasis, we evaluated the germinal center (GC) IgA responses in NCD- and HFHC-fed mice. Notably, we discovered that the frequency of GL7^+^ CD95^+^ GC B cells was substantially reduced in the PP of HFHC-fed mice, compared with NCD controls (Figure 2A-B). As a result, the differentiation of IgA^+^ B cells in the PPs was also diminished (Figure 2C-D), suggesting that NASH induces a defective GC response and a loss of IgA-producing cells. To confirm these findings, we performed immunofluorescence staining of PPs from NCD- and HFHC-fed mice and observed smaller GC regions in the PP of NASH mice (Figure 2E). We then used histocytometry to analyze the immunofluorescence data and found a reduction in GC B cell responses in NASH mice (Figure 2E-F), in agreement with our earlier observations. Given that GCs play a crucial role in the affinity maturation of IgA, we reasoned that impaired GC B cell responses could affect the ability of IgA to bind commensal bacteria. Indeed, we detected a lower IgA coating of fecal bacteria in NASH mice (Figures 2G-I). Together, these data suggest that HFHC diet-induced NASH has profound detrimental effects on intestinal immune responses, including a reduced GC reaction and impaired IgA production and ability to bind bacteria.

**Figure 2.**
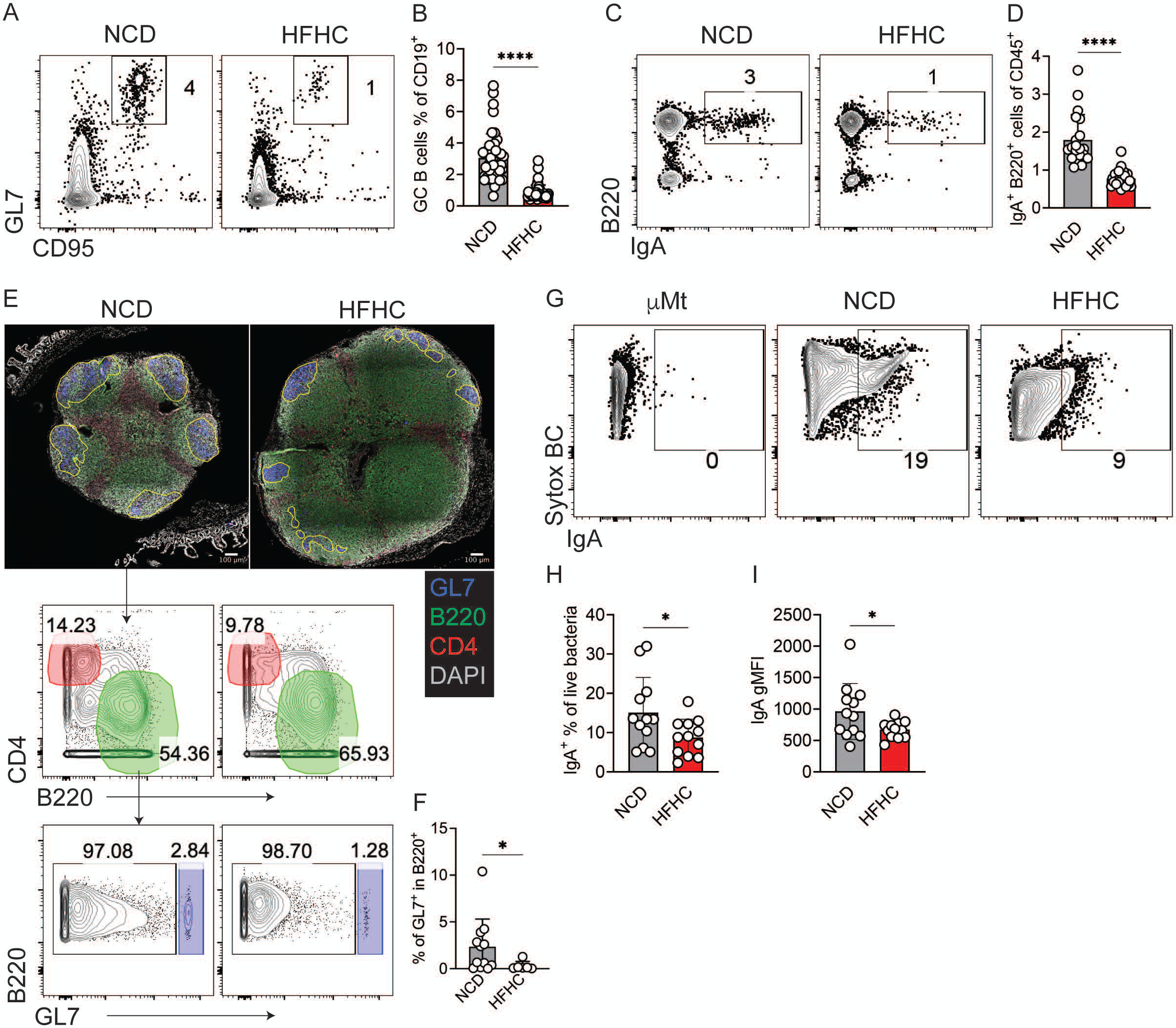
Impaired GC responses and decreased IgA+ B cells in HFHC mice. (A) Representative flow cytometry analysis of GC (CD95^+^ GL7^+^) among total CD19^+^ B cells, in PPs of NCD and HFHC mice. (B) Frequency of GC among total CD19^+^ B cells. (C) Representative flow cytometry analysis of IgA^+^ B cells (B220^+^ IgA^+^) among total CD45^+^ cells, in PPs of NCD and HFHC mice. (D) Frequency of IgA^+^ B cells among total CD45^+^ cells. (E) Immunofluorescent staining of PPs from NCD and HFHC mice (Top), GL7 (blue), B220 (green), CD4 (red), DAPI (grey). Histocytometric analysis of B220 and CD4 expression from the immunofluorescent staining (Middle). Histocytometric analysis of GC (B220^+^ GL7^+^) among CD4^−^ B220^+^ cells (Bottom). (F) Frequency of GC among CD4^−^ B220^+^ cells. (G) Representative flow cytometry analysis of IgA coating of fecal bacterial, live bacterial was gated as DAPI^−^ Sytox BC^+^. (H) Frequency of IgA coated bacterial among total live bacterial. (I) Geometric mean fluorescence intensity (gMFI) of IgA in the IgA^+^ bacterial. Data are representative of three independent experiment, ns=not significant, ****p<0.001, *p<0.05, unpaired t test.

### Dysfunctional Intestinal Tfh Cells in HFHC Diet-Induced NASH Mice

We next sought to functionally characterize the Tfh cells in the PPs of HFHC diet-induced NASH mice. Flow cytometry analysis revealed an altered Tfh cell phenotype in PP from NASH mice, including enhanced CD40L and CD44 expression but a substantially decreased expression of PD1 and BCL6 (Figure 3A-B). Bulk RNA-Seq of FACS-sorted Tfh cells from the PP of NCD and HFHC-fed mice revealed a marked shift in the transcriptional landscape of Tfh cells in NASH (Figure 3C). We found that genes that were highly expressed in Tfh cells from the PP of NCd mice were markedly downregulated in NASH (Figure 3C). To better understand the nature of this shift, we performed Gene Set Enrichment Analysis (GSEA) on the differentially expressed genes and discovered that genes involved in the differentiation of Tfh cells were downregulated in the PP of HFHC-fed mice, suggesting a loss of Tfh cell identity during NASH (Figure 3D). Specifically, we detected a lower expression of key Tfh cell signature genes and transcription factors that are vital for Tfh cell differentiation and functionality such as *Bcl6*, *S1pr2*, and *Pdcd1* (Figure 3E). Tfh cells from NASH mice also showed a substantial reduction in IL-4 gene expression (Figure 3E) and production in vivo, detected using IL-4 GFP mice (Supplementary Figure 2A-B). In addition, NASH Tfh cells showed no changes in the Th1 transcription factor T-bet but a decreased expression of GATA3 (Th2) and increased ROR-γt, master regulator of the Th17 fate (Figure 3F and Supplementary Figure 2C-D,). Tfh cells from NASH mice had increased expression of the Tfh cell differentiation inhibitor KLF2^27^ (Figure 3F) and its upstream regulator FOXO1^28^ (Figure 3F-G). Chemokine receptors such as CCR7 and S1PR1 were also increased in Tfh cells from the PP of NASH mice (Figure 3E, H), suggesting a dysfunctional migration. Furthermore, Tfh cells from NASH mice had increased glucose uptake (Supplementary Figure 2E), decreased mitochondria mass (Supplementary Figure 2F), and lower mitochondria membrane potential (Supplementary Figure 2G), indicative of altered metabolism.

**Figure 3.**
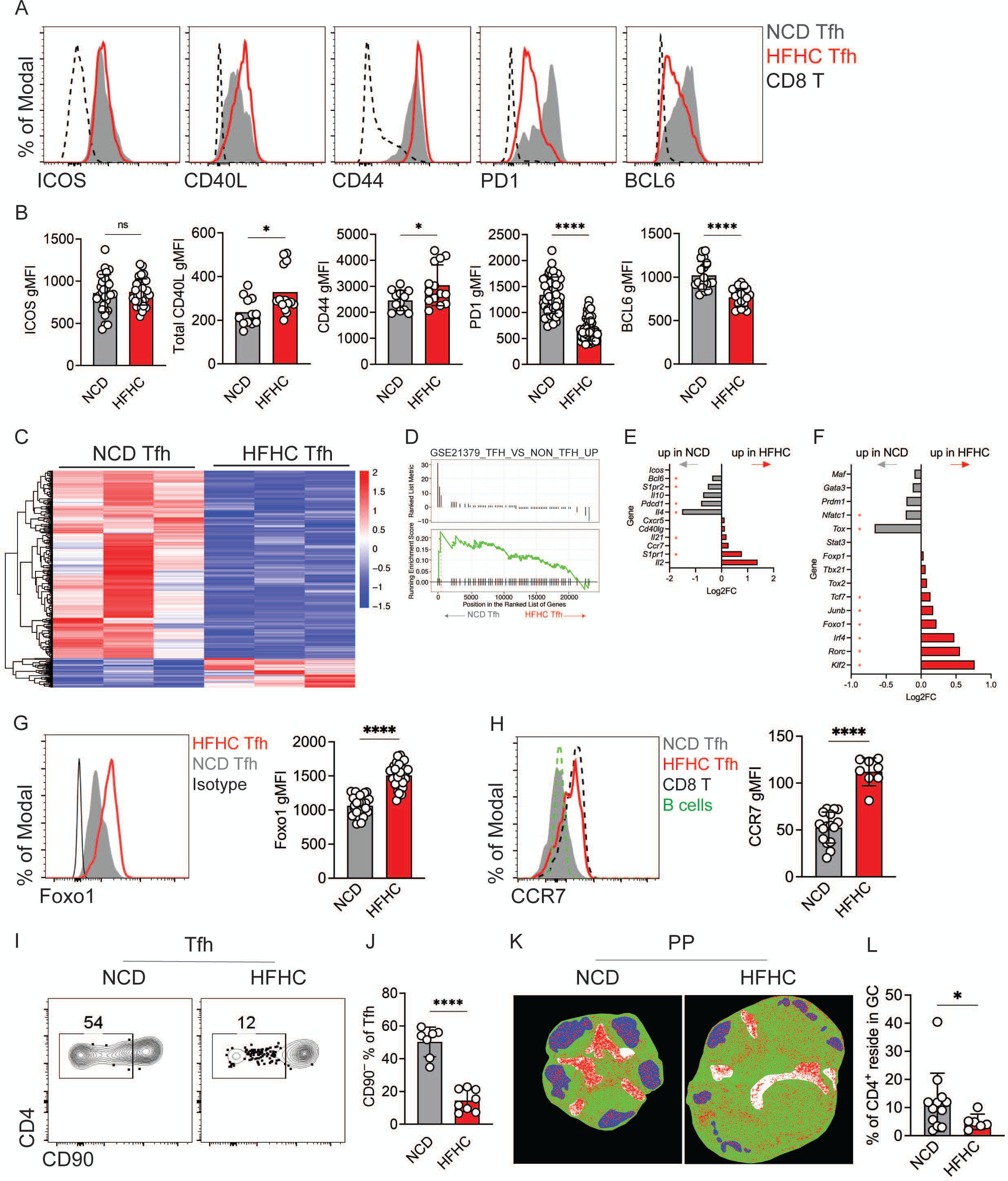
Disturbed phenotype of intestinal Tfh cells in HFHC mice. A) Representative flow cytometry histogram of ICOS, CD40L, CD44, PD1, BCL6 expression in NCD Tfh (grey), HFHC Tfh (red) and CD8 T (black) cells in PPs. (B) gMFI of ICOS, CD40L, CD44, PD1, BCL6 in NCD Tfh (grey) and HFHC Tfh (red) cells in PPs. (C) Heatmap of differentially expressed genes between NCD and HFHC Tfh cells in bulk RNA-Seq data (fold change>2, false discovery rate<0.05). (D) GSEA plot of ranked genes in NCD Tfh vs HFHC Tfh cells identified in bulk RNA-Seq of sorted Tfh cells to specific gene set (GSE21379_TFH_VS_NON_TFH_UP). (E) Change of Tfh signature genes between NCD and HFHC Tfh cells. (F) Change of transcription factor genes between NCD and HFHC Tfh cells. (G) Representative flow cytometry histogram of FOXO1 expression in HFHC Tfh and NCD Tfh cells (Left), gMFI of FOXO1 in HFHC Tfh and NCD Tfh cells (Right). (H) Representative flow cytometry histogram of CCR7 expression in HFHC Tfh, NCD Tfh, CD8 T and B cells (Left), gMFI of CCR7 in HFHC Tfh and NCD Tfh cells (Right). (I) Representative flow cytometry analysis of CD90 expression in HFHC Tfh and NCD Tfh cells. (J) Frequency of CD90^low^ Tfh cells in PPs of NCD and HFHC mice. (K) Histocytometry analysis of the localization of CD4 T cells in PPs of NCD and HFHC mice. (L) Frequency of CD4 T cells reside in the GCs of PPs in NCD and HFHC mice by histocytometry analysis. Data are representative of three independent experiment, ns=not significant, ****p<0.001, *p<0.05, unpaired t test.

In a properly organized germinal center (GC), Tfh cells residency within the GC is facilitated by the downregulation of KLF2, CCR7, and S1PR1^27^. The increase in these molecules in Tfh cells from NASH mice led us to interrogate their localization within the GCs, an important aspect needed for maintaining a robust GC response and B cell maturation^29^. A recent study showed that primary Tfh cells include a subset of CD90^low^ GC-resident and CD90^+^ non-resident cells^29^. Thus, we investigated how NASH affects the location of the Tfh cells in the GC and found that, unlike Tfh cells from NCD mice, the majority of Tfh cells in the PP of NASH mice were CD90^+^ non-resident GC cells while the CD90^low^ resident cells decreased (Figure 3I-J). Indeed, histocytometry analysis of PPs showed fewer CD4 T cells within the GCs of NASH mice, compared with controls (Figure 3K-L). Thus, these data show that Tfh cells present with a dysfunctional phenotype that includes a defective differentiation transcriptional program, decreased IL4 production, and a loss of residence within the GC.

Considering the potential contribution of obesity per se to a dysfunctional Tfh phenotype in NASH mice, we explored the potential impact of genetic obesity on PP Tfh cells. To this end, we utilized Ob/Ob (*Lep*^ob^) mice that develop hyperphagia, obesity, intestinal bacterial overgrowth, and liver steatosis despite being fed an NCD. By 25 weeks of age, the NCD-fed Ob/Ob mice gained substantially more bodyweight than their WT counterparts (Supplementary Figure 3A). Despite such profound obesity, however, we observed no substantial alterations in the frequency, GC-residency, and expression of PD1, BCL6, and Foxo1 in the Tfh cells of Ob/Ob mice (Supplementary Figure 3B-F). Furthermore, while there was no reduction in the GC reaction (Supplementary Figure 3G), there was a slight decrease in the frequency IgA^+^ B cells (Supplementary Figure 3H) in the PP of Ob/Ob mice. This data suggests that the Tfh cell dysfunction caused by NASH may not be directly attributable to obesity but to the effects of the HFHC diet and/or subsequent NASH progression.

### Microbial Unresponsiveness of Intestinal Tfh Cells in Diet-Induced NASH Mice

IgA responses, especially those mediated by high-affinity T cell-dependent IgA, play a crucial role in maintaining intestinal homeostasis, regulating microbiota community, bolstering mucosal defense, and restraining invasive commensal species^30–32^. The production of intestinal IgA is primarily attributed to the interactions between Tfh cells and B cells within the GC in PPs^14^. Given that intestinal Tfh-GC responses can be induced and modulated by gut microbiota^19, 20^, we sought to determine whether Tfh cells in NASH remain responsive to gut microbiota. To accomplish this, we depleted the gut microbiota of NCD- and HFHC-fed mice with broad-spectrum antibiotics provided in the drinking water (Supplementary Figure 4A). The antibiotic treatment effectively depleted the gut bacteria and altered the microbiota diversity and composition in both NCD and HFHC mice (Supplementary Figure 4B-E). Notably, antibiotic treatment resulted in a substantial reduction in the frequency of Tfh cells, GC B cells, and IgA-producing cells in PPs of NCD-, but not HFHC-fed mice (Supplementary Figure 4F-G). Similarly, antibiotic treatment altered the expression of PD1 and Foxo1 in Tfh cells from NCD but not HFHC mice (Supplementary Figure 4H), suggesting that Tfh cells are unresponsive to changes in the gut microbiota during NASH. One possibility is that the lack of dependency of the NASH Tfh cells on the microbiota is related to a decreased T cell receptor (TCR) signal in the pre-Tfh subset (Supplementary Figure 4I).

### Tfh Cells Maintain Intestinal Barrier and Protect against NASH

To investigate the direct role of Tfh cells in NASH progression, we used *Cd4*^Cre^ *Bcl6*^flox/flox^ mice that specifically lack Tfh cells (Figure 4A). Both Tfh knockout (Tfh KO) and littermate wild-type (Wt) mice were fed the HFHC diet for up to 20 weeks to induce NASH. The absence of Tfh cells led to the abrogation of GCs and IgA^+^ B cells in PPs (Figure 4B, Supplementary Figure 5A), along with a notable decrease in IgA^+^ B cells in the small intestine lamina propria (sLPL) (Supplementary Figure 5B). Given the critical role of intestinal immune cells in maintaining gut homeostasis and modulating microbiota composition^14–16, 18^, we performed 16S rRNA sequencing to analyze microbiota communities of Tfh KO and Wt mice. Our analysis revealed that the microbiota communities were substantially distinct between Tfh KO and Wt mice as failed to cluster by principal component analysis (PCA) (Figure 4C). Using targeted mass spectrometry, we also examined microbiota-derived short-chain fatty acids (SCFA) and found a substantial reduction of butyrate in fecal contents from Tfh KO microbiota (Figure 4D). Given the importance of butyrate in maintaining intestinal integrity^12^, we assessed the levels of the tight junction proteins in intestinal epithelial cells and found that αE-Catenin and Occludin were reduced in Tfh KO mice, compared with Wt controls (Figure 4E-F). As a result, we observed a higher intestinal permeability in Tfh KO mice following a FITC-Dextran oral gavage assay (Figure 4G). In the PPs and small intestine LPL, we detected an increased accumulation of pro-inflammatory CD44^+^ T-bet^+^ CD8 T cells and a decrease in regulatory T cells (Tregs) in the mLN (Supplementary Figure 5C-G), a finding consistent with the requirement for butyrate for the differentiation of Tregs ^33, 34^. Together, these results suggest a disrupted gut homeostasis and the development of a “leaky gut” in the absence of Tfh cells.

**Figure 4.**
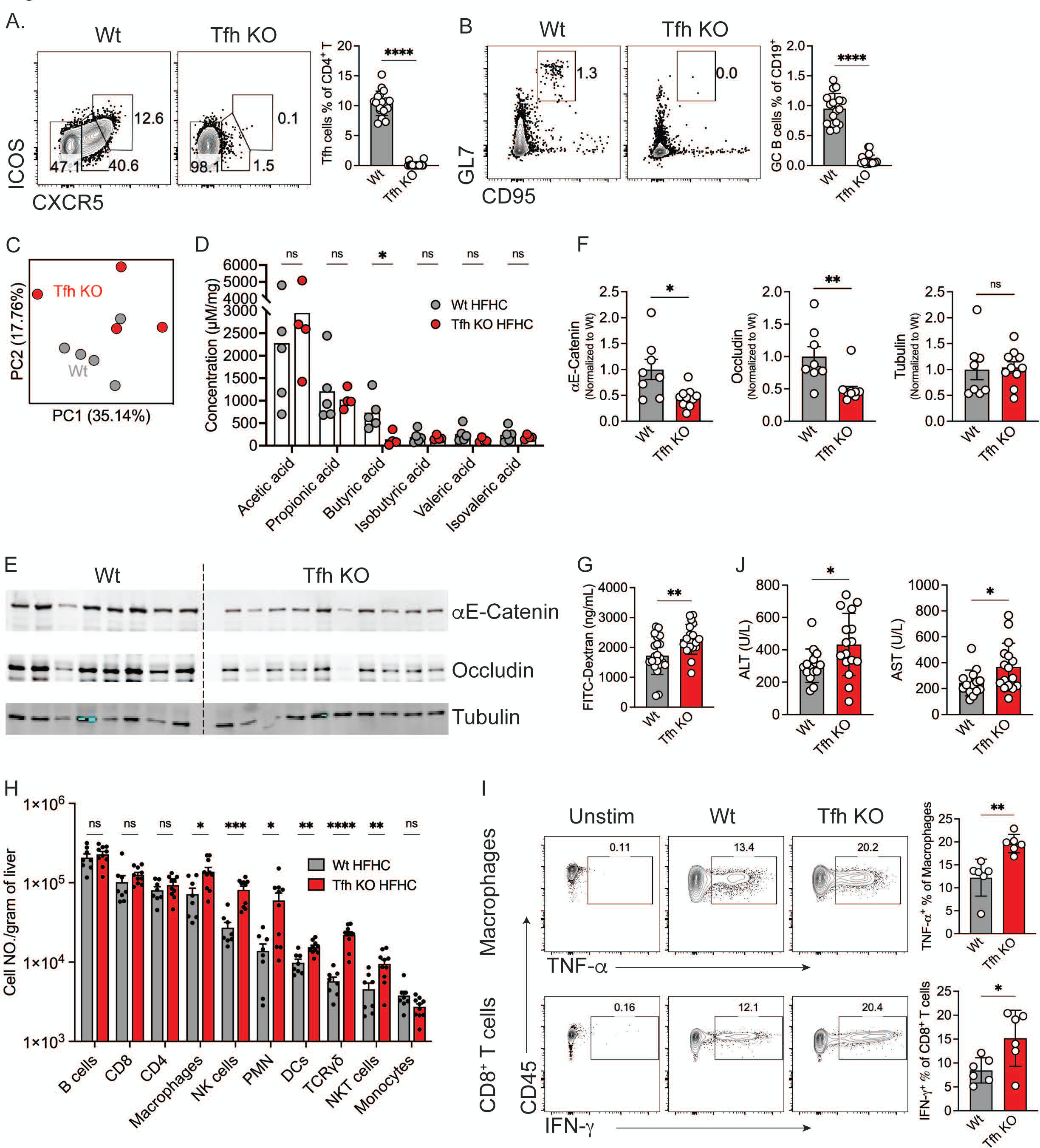
The role Tfh cells in gut homeostasis and NASH progression. (A) Representative flow cytometry analysis to identify Tfh (CXCR5^high^ ICOS^high^), Pre-Tfh (CXCR5^intermediate^ ICOS^intermediate^) and Non Tfh (CXCR5^−^ ICOS^−^) cells among Foxp3^−^ CD4 T cells, in PPs of Wt and Tfh KO mice fed on HFHC diet (Left), frequency of Tfh cells among total CD4 T cells (Right). (B) Representative flow cytometry analysis of GC (CD95^+^ GL7^+^) among total CD19^+^ B cells, in PPs of Wt and Tfh KO mice fed on HFHC diet (Left), frequency of GC cells among total CD19+ B cells (Right). (C) PCA plot of 16S rRNA sequencing of intestinal microbiota from Wt and Tfh KO mice. (D) Quantification of short-chain fatty-acids in fecal pellets from Wt and Tfh KO mice by targeted mass spectrometry. (E) Western blot of tight junction proteins (Occludin, αE-Catenin) and housekeeping protein (Tubulin) in intestinal epithelial cells of Wt and Tfh KO mice. (F) Quantification of tight junction proteins (αE-Catenin, right; Occludin, middle) and housekeeping protein (Tubulin, left) from western blot. (G) Concentration of FITC-Dextran in blood of Wt and Tfh KO mice 4 hours after FITC-Dextran oral gavage assay. (H) Cell number of different immune cells identified by CyTOF in the liver of Wt and Tfh KO mice. (I) Representative flow cytometry analysis of cytokines production (TNF-α, IFN-γ) by ex vivo macrophages (Left top) and CD8 T cells (Left bottom) after stimulation with LPS. Frequency of TNF-α+ macrophages (Right top) and IFN-γ+ CD8 T cells (Right bottom) among total macrophages and CD8 T cells, respectively. (J) ALT (Right) and AST (Left) level in serum of Wt and Tfh KO mice. Data are representative of three independent experiment, ns=not significant, ****p<0.001, ***p<0.005, **p<0.01, *p<0.05, unpaired t test or ANOVA.

Since disruption of the intestinal barrier is known to exacerbate liver inflammation in NASH^6, 8, 35^, we examined the immune infiltrates in the liver of HFHC-fed Tfh KO mice. Comprehensive profiling of the immune landscape of the liver using cytometry by time of flight (CyTOF) showed an overall increased immune cell infiltration in Tfh KO mice (Figure 4H), suggesting exacerbated inflammation. Notably, hepatic macrophages and CD8 T cells from Tfh KO mice produced increased levels of TNF-α and IFN-γ, respectively (Figure 4I). As a result of increased inflammation, the circulatory levels of liver function enzymes alanine transaminase (ALT) and aspartate transaminase (AST) were higher in Tfh KO mice, compared with Wt controls (Figure 4J). In conclusion, our data indicate that the protective role of intestinal Tfh cells in the gut mitigate liver inflammation and disease progression in NASH.

### KLF2 Insufficiency Improves the Tfh Cell Phenotype and Ameliorates NASH

Given that Tfh cells are crucial for maintaining gut and liver immune homeostasis and that they present a dysfunctional phenotype in NASH, we reasoned that restoring the function of Tfh cells in NASH may provide protection against the disease. In our RNAseq analysis, we found an abnormal increase in KLF2 in Tfh cells from the PP of NASH mice likely due to the role of KLF2 in restricting the fate commitment of Tfh cells^27^. Therefore, we hypothesized that reducing KLF2 in Tfh cells during NASH would restore their differentiation and function and may ameliorate disease progression. To decrease KLF2 expression in Tfh cells, we used *Cd4*^Cre^ *Klf2*^Wt/flox^ (KLF2^Δ/+^) mice that harbor a single allele deletion of the *Klf2* gene in CD4 T cells. We fed KLF2^Δ/+^ and littermate Wt mice a HFHC diet to induce NASH and assessed the phenotype of Tfh cells in the PPs and NASH severity. Compared with Wt controls, KLF2^Δ/+^ mice showed no alterations in the frequency of total Tfh cells, although a slight increase in pre-Tfh cells was observed (Figure 5A-B). Importantly, Tfh cells in KLF2^Δ/+^ mice had an improved phenotype as evidenced by an increased expression of BCL6 and PD1 (Figure 5C-D), while FOXO1 and ICOS expression remained unchanged (Figure 5E). Importantly, the restored Tfh cell phenotype in HFHC-fed KLF2^Δ/+^ mice correlated with an increase in IgA^+^ B cells (Figure 5F) and overall ameliorated NASH. Specifically, KLF2^Δ/+^ mice had lower liver weight (Figure 5G), reduced triglycerides (Figure 5H), and a decreased NAFLD activity score (NAS) determined in H&E-stained of liver sections (Figure 5I). Furthermore, KLF2^Δ/+^ mice had lower ALT and AST levels (Figure 5J). Consistent with the improvements in NASH, KLF2^Δ/+^ mice displayed improved glucose and insulin tolerance (Supplementary Figure 6 A-B) and energy metabolism as assessed by indirect calorimetry (Supplementary Figure 6C). We also performed bulk RNA-Seq of the whole liver tissue and conducted GSEA for a set of NASH signature genes and found a marked downregulation of NASH-associated genes in KLF2^Δ/+^ mice, compared with Wt controls (Figure 5K). Moreover, KLF2^Δ/+^ mice had a decreased hepatic accumulation of pathogenic PD1^+^ CD8 T cells^36^ (Figure 5L). Overall, our findings underscore a key role for KLF2 in regulating Tfh cell function and NASH pathogenesis.

**Figure 5.**
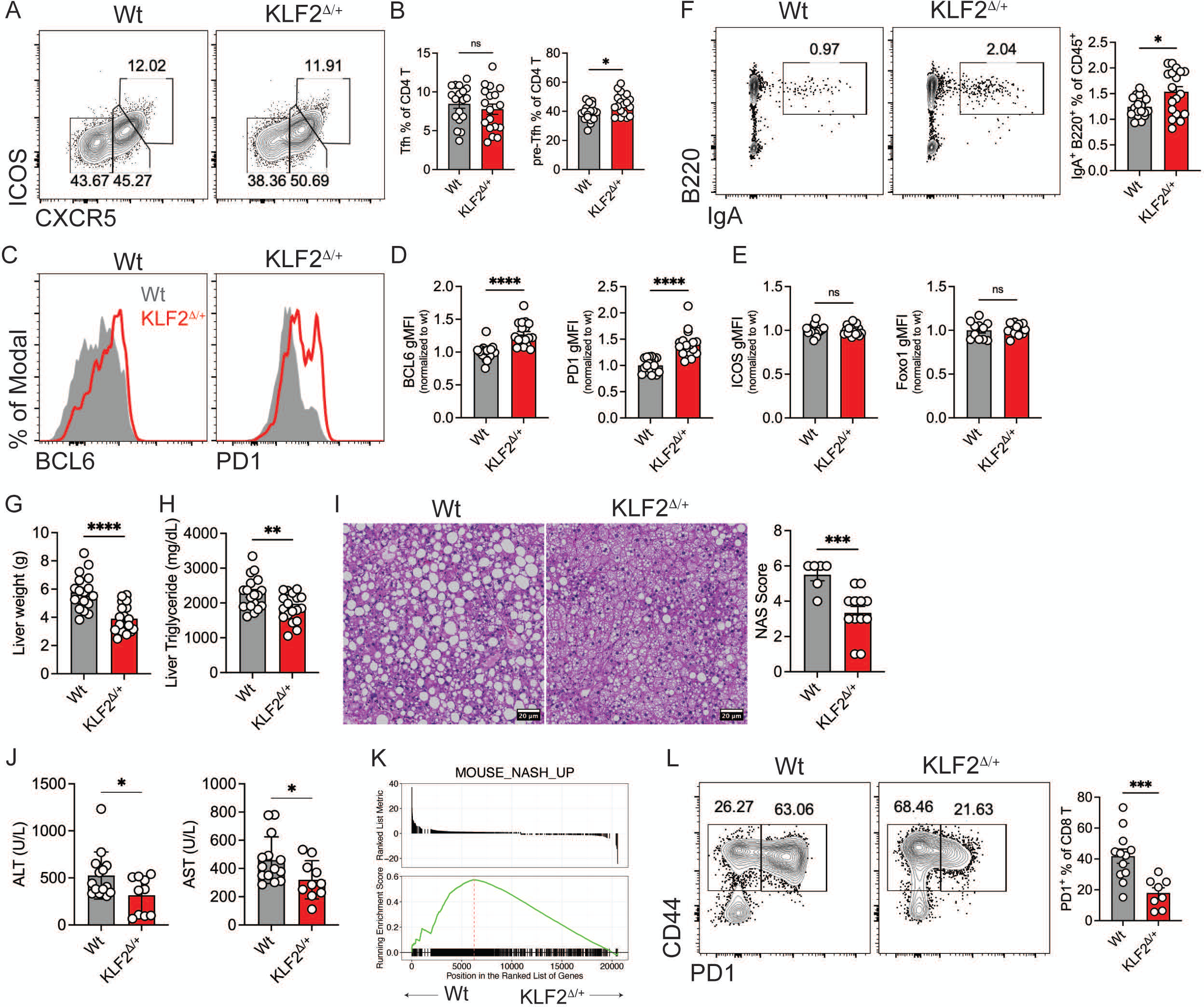
Genetic reduction of KLF2 improves Tfh cell phenotype and mitigates NASH pathogenesis. (A) Representative flow cytometry analysis to identify Tfh (CXCR5^high^ ICOS^high^) and Pre-Tfh cells (CXCR5^intermediate^ ICOS^intermediate^) among Foxp3^−^ CD4 T cells in PPs of Wt and KLF2^τι/+^ mice. (B) Frequency of Tfh (Left) and Pre-Tfh cells (Right) among total CD4 T cells. (C) Representative flow cytometry histogram of BCL6 (Left) and PD1 (Right) expression Tfh cells from Wt and KLF2^Δ/+^ mice. (D) gMFI of BCL6 (Left) and PD1 (Right) in Tfh cells from Wt and KLF2^Δ/+^ mice. (E) gMFI of ICOS (Left) and FOXO1 (Right) in Tfh cells from Wt and KLF2^Δ/+^ mice. (F) Representative flow cytometry analysis to identify IgA^+^ B cells (IgA^+^ B220^+^) among CD19^+^ B cells (Left), frequency of IgA^+^ B cells among CD19^+^ B cells. (G) Liver weight of Wt and KLF2^Δ/+^ mice. (H) Quantification of triglyceride in liver tissue. (I) Representative H&E staining of liver section (Left), NAS score from the H&E staining (Right). (J) ALT (Right) and AST (Left) level in serum of Wt and KLF2^Δ/+^ mice. (K) GSEA plot of ranked genes in Wt vs KLF2^Δ/+^ mice identified in bulk RNA-Seq of liver tissues to MOUSE_NASH_UP gene set generated by our own bulk RNA-Seq of liver tissue from NCD and HFHC mice. (L) Representative flow cytometry analysis to identify PD1^+^ CD8 T cells (CD44^+^ PD1^+^) among total CD8 T cells (Left), frequency of PD1^+^ CD8 T cells among total CD8 T cells (Right). Data are representative of three independent experiment, ns=not significant, ****p<0.001, ***p<0.005, **p<0.01, *p<0.05, unpaired t test.

## DISCUSSION

The gut-liver axis involves bidirectional communication between the gut and liver and its dysfunction contributes to the onset and progression of NAFLD^6, 8^. In addition, dysbiosis of the gut microbiota can influence energy homeostasis, lipid metabolism, and fat storage, leading to worsened liver disease^7, 8^. Mechanistically, an increased permeability of the gut epithelium or “leaky gut” caused by factors such as diet, pathogens, or chemicals, facilitates the translocation of microbial antigens to the liver^7, 9^. Lipopolysaccharides (LPS) are the main gut-derived antigen that can induce metabolic endotoxemia and promote hepatic steatosis, inflammation, and fibrosis^7, 9^. Additionally, LPS directly activates innate immune responses in the liver, resulting in an inflammatory process that contributes to tissue damage^5, 7, 9^. Thus, a better understanding of the immune mechanisms regulating the inflammatory tone in the gut-liver axis is needed for the development of effective therapeutic strategies against NASH.

Tfh cells play a pivotal role in the adaptive immune responses to pathogens^37, 38^. These cells are primarily found within the follicles of secondary lymphoid organs, including the lymph nodes and the spleen, where they participate in the formation of GCs that are critical for the production of high-affinity antibodies^37, 38^. Notably, Tfh cells have emerged as important regulators of gut homeostasis^14, 16^. In our study, we found that NASH induces substantial alterations in the cellularity and gene expression of PPs including an accumulation of Tfh cells. The expansion of Tfh cells, however, was accompanied by a dysregulated Tfh phenotype, reduced IgA+ B cell differentiation, and diminished GC responses. Such dysfunction of Tfh cells during NASH is likely responsible for the abnormal GC and IgA responses as demonstrated by our experiments with Tfh-deficient mice. Moreover, we found that NASH upregulated the Tfh expression of KLF2, a transcription factor that restricts the fate commitment of these cells. Overall, the atypical expression pattern of Tfh genes suggests a loss of Tfh cell identity in NASH, which could be a major contributing factor to the impaired GC responses and decreased IgA production. Despite our findings, the molecular mechanisms contributing to the upregulation of KLF2 in Tfh cells during NASH remain elusive. Previous studies demonstrated that ICOS signaling downregulates KLF2 expression T cells, which is a critical step for Tfh cell differentiation. However, we did not detect changes in ICOS expression in Tfh cells during NASH. This finding suggests the involvement of additional factors that regulate KLF2 expression in NASH.

Interestingly, our results showed that the dysfunction of Tfh cells is not directly attributable to obesity but is specific to the effects of HFHC diet and/or NASH development, as suggested by the normal Tfh cell phenotype and function in the Ob/Ob (*Lep*^ob^) mouse in which obesity is largely attributed to their leptin deficiency-induced hyperphagia^39^. Given this surprising finding, we speculate that the HFHC diet, rather than obesity per se, may be the primary factor triggering Tfh cell dysfunction. Indeed, recent research showed that dietary sugar, high in our HFHC diet, disrupts the protective role of the immune system against metabolic disease^40^. Future work is needed to isolate specific dietary components responsible for this effect and to identify the mechanisms by which they disrupt intestinal immune homeostasis.

One of the intriguing findings of our work is the unresponsiveness of Tfh cells to the gut microbiota changes induced by NASH. It is well established that gut microbiota can initiate and fine-tune the Tfh-GC interactions^32, 41^. However, our observations indicate that Tfh cells in NASH mice lose this regulatory control, perpetuating defective GC responses and sub-optimal IgA production even after alterations in the microbiota. Such decoupling of Tfh function from changes in the gut microbiota may contribute to NASH progression. However, the mechanisms leading to the unresponsiveness of Tfh cells to the microbiota in NASH remain to be elucidated. Our data showed a significant upregulation of KLF2 in NASH Tfh cells, which increased the expression of chemokine receptors such as CCR7 and S1PR1. Notably, CCR7 guides T cell localization to T cell zones within lymphoid tissues, while S1PR1 governs T cell trafficking and re-entry into circulation. Under homeostasis, the differentiation of Tfh cells is facilitated by a tightly controlled shift in chemokine receptor expression whereby pre-Tfh cells downregulate CCR7 and upregulate CXCR5, facilitating their migration from T cell zones to B cell follicles to assist in GC formation^38^. Given the altered expression of chemokine receptors in NASH Tfh cells, it is plausible that these cells might be mislocalized, which may affect their function and responsiveness to microbial antigens. Supporting this notion, our histocytometry analyses revealed an aberrant dispersion of intestinal Tfh cells within PPs in NASH mice. Additionally, the antigen specificity of Tfh cells, determined by their TCR, may be altered in NASH as evidenced by the loss of TCR signal in pre-Tfh cells. TCR sequencing of Tfh cells from Nin NASASH mice is needed to confirm this possibility.

In summary, our study highlights the complex interplay between intestinal Tfh cell function and NASH. The paradoxical expansion of dysfunctional Tfh cells in NASH leads to sub-optimal GC responses, reduced IgA production, and disturbed gut homeostasis. Therefore, we propose that strategies aimed at restoring Tfh cell function, such as downregulation of KLF2, could offer a viable immune therapy for the treatment of NASH.

## EXPERIMENTAL PROCEDURES

### Experimental Animals

Wild-type (Wt) C57BL/6J, *Cd4*^Cre^ (B6.Cg-Tg(Cd4-cre)1Cwi/BfluJ), *Bcl6*^flox/flox^ (B6.129S(FVB)-Bcl6tm1.1Dent/J), Ob/Ob (B6.Cg-Lepob/J) mice were purchased from The Jackson Laboratory. *Nur77*^GFP^ mice on B6 background were kindly provided by Dr. Kristin A. Hogquist at the University of Minnesota. *Kfl2*^flox/flox^ mice on B6 background were kindly provided by Dr. Stephen C. Jameson at the University of Minnesota. At 6 weeks of age, mice received a normal chow diet (NCD) or high-fat high-carbohydrate diet (HFHC) (40%kcal palm oil, 20%kcal fructose, and 2% cholesterol) supplemented with 42g/L of carbohydrates in the drinking water (55% Fructose, 45% Sucrose; Sigma) ^42^. All mice were male, age-matched, and housed in a standard pathogen-free environment. Animal experiments followed the Guide for the Care and Use of Laboratory Animals and approved by the University of Minnesota Institutional Animal Care and Use Committee.

### Immune Cell Isolation and Characterization

Immune cells were isolated from the colon lamina propria as previously described ^43^. In cytometry by time-of-flight (CyTOF) analysis, cells were stained with 0.5 µg of metal-conjugated primary antibodies (Fluidigm) for 30 mins at 4°C. Data was acquired on a CyTOF2 (DVS Sciences) and analyzed using Cytobank. For flow cytometry, cells were incubated with fluorophore-conjugated primary antibodies for 30 mins at 4°C. For intracellular staining of transcription factors, cells were stained with antibodies to surface makers, then fixed and permealized with True-Nuclear™ Transcription Factor Buffer Set (Biolegend) and were incubated with antibodies to T-bet (4B10, Biolegend), ROR-γt (Q31-378, BD Biosciences), Foxp3 (150D, Biolegend). Viability dye LIVE/DEAD™ Fixable Near IR (780) Viability Kit was from Thermofisher. The list of other antibodies used is as follows and were purchased from eBioscience, BD Biosciences or BioLegend, unless otherwise indicated: anti-CD4 (GK1.5 or RM4-4), CD8α (53–6.7), anti-CD24 (M1/69), anti-NK1.1 (PK136), anti-CD44 (IM7), anti-TCRβ (H57-597), anti-PD-1 (29F.1A12), anti-CD45.1 (A20), anti-CD45.2 (104), anti-B220 (RA3-6B2), anti-CD11c (N418), anti-CD11b (M1/70), anti-F4/80 (BM8). Flow cytometry data were acquired on a BD Fortessa (BD Biosciences) and analyzed using Flowjo software.

### Bulk RNA sequencing

Total RNA was extracted from livers using an RNeasy Plus Mini kit (Qiagen). Samples were sequenced on a Novaseq 6000 using a 150 PE flow cell at the University of Minnesota Genomics Center. The SMARTer Stranded RNA Pico Mammalian V2 kit (Takara Bio) was used to create Illumina sequencing libraries. Differential gene expression analysis was performed using edgeR (Bioconductor). Gene set enrichment analysis (GSEA) was performed using Clusterprofiler (Bioconductor).

### Histocytometry

The histocytometry analysis was described previously ^44–46^. Briefly, the region of interests (ROIs) was identified and the fluorochrome intensities of each ROI were quantified using ImageJ and data were exported into Excel, Prism and FlowJo software for the localization analysis.

### Histology

Liver tissues fixed in 10% formalin were histologically assessed for steatosis by hematoxylin and eosin staining, analysis of NAFLD activity score (NAS) was performed by a blinded liver pathologist.

### Metabolic assessments

Insulin tolerance tests (ITT) and glucose tolerance tests (GTT) were performed as previously described ^5^. To determine energy metabolism, mice were placed in automated metabolic cages for 48 h. An assessment of energy expenditure (EE) was obtained via indirect calorimetry in free-moving animals housed in individual cages consisting of an indirect open circuit calorimeter that provides measures of O2 consumption and CO2 production (Oxymax, Columbus Instruments, Ohio). The ambulatory activity was assessed by the breaking of infrared laser beams in the x-y plane. The cages were provided with ad libitum access to food and water throughout the procedure. These procedures were performed by the IBP Phenotyping Core at the University of Minnesota.

### Statistical Analysis

Statistical significance between means was determined with an unpaired t-test using GraphPad Prism 8.3. Spearman’s rank test was used to determine correlation. Data are presented as means ± SEM. Statistical significance was set to 5% and denoted by *p < 0.05, **p < 0.01, and *** p < 0.001.

## ACKNOWLEDGMENTS

This study was supported by the National Institute of Diabetes and Digestive and Kidney Diseases (DK122056 to X.S.R.) and the National Heart, Lung, and Blood Institute (HL155993 to X.S.R.). G.F. is supported by a NIH training grant (T32DK083250). We recognize the staff from the Research Animal Resources, University Flow Cytometry Resource, Center for Metabolomics and Proteomics, Genomics Center, and the Clinical and Translational Science Institute at the University of Minnesota for their assistance.

## FIGURE LEGENDS

**Supplementary Figure 1 (related to Figure 1).**
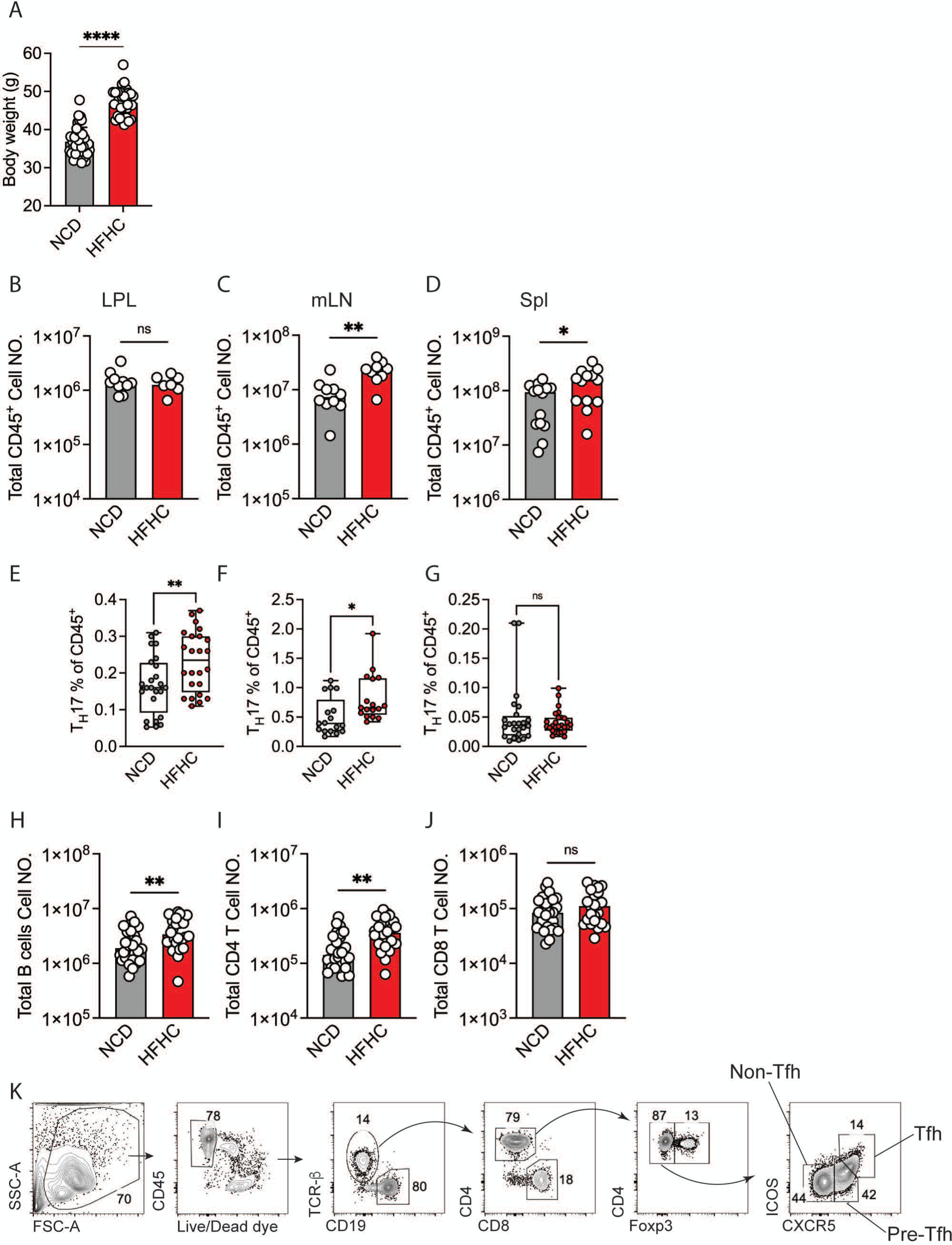
(A) Body weight of NCD and HFHC mice. (B) Cell number of total live CD45^+^ cells in small intestine LPL. (C) Cell number of total live CD45^+^ cells in mLN. (D) Cell number of total live CD45^+^ cells in spleen. (E) Frequency of Th17 cells among total live CD45^+^ cells in small intestine LPL. (F) Frequency of Th17 cells among total live CD45^+^ cells in mLN. (G) Frequency of Th17 cells among total live CD45^+^ cells in spleen. (H) Cell number of total B cells in PPs. (I) Cell number of total CD4 T cells in PPs. (J) Cell number of total CD8 T cells in PPs. (K) Representative flow cytometry gating strategy for identifying Tfh cells in PPs. Data are representative of three independent experiment, ns=not significant, ****p<0.001, **p<0.01, *p<0.05, unpaired t test.

**Supplementary Figure 2 (related to Figure 3).**
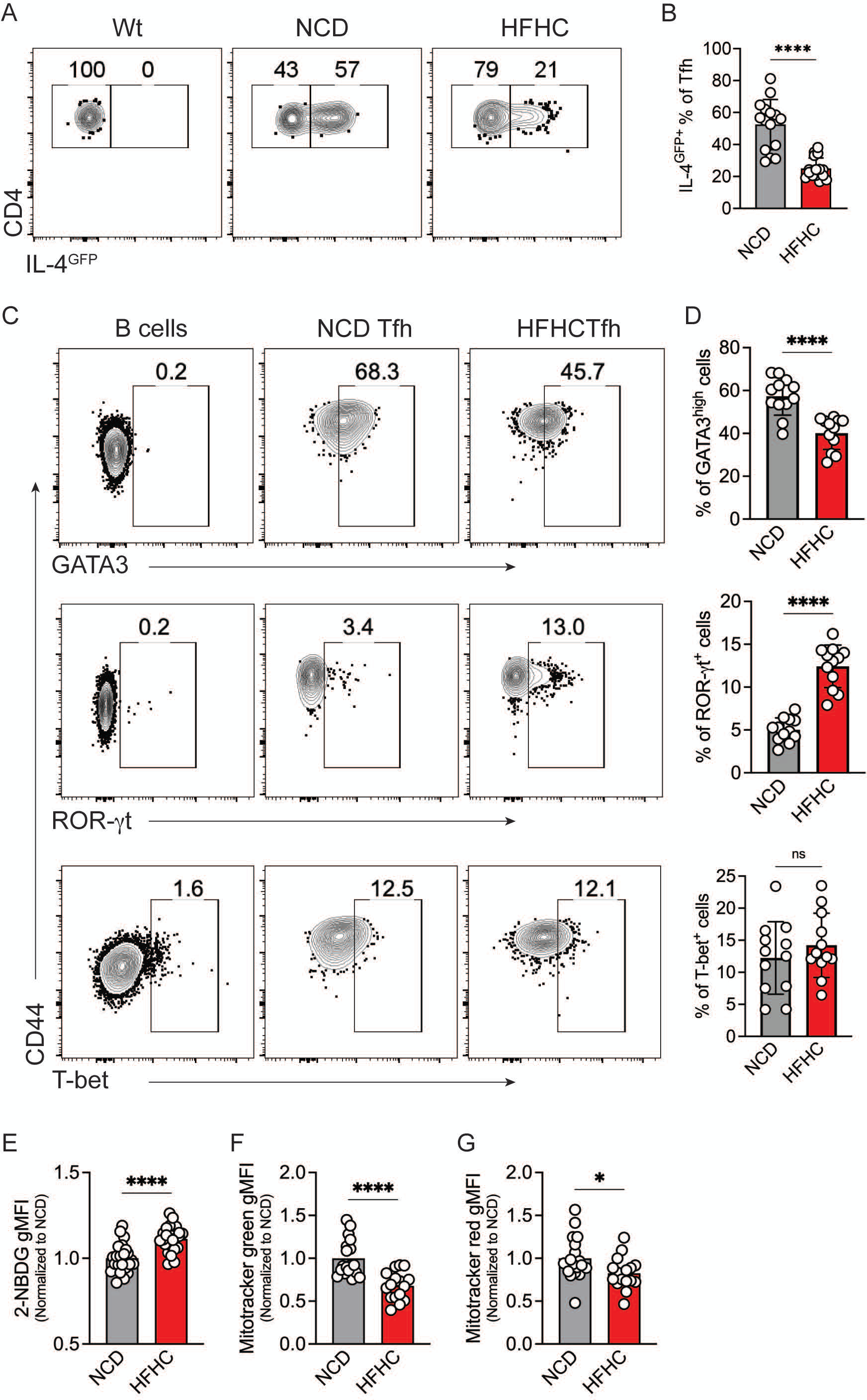
(A) Flow cytometry analysis to identify IL-4^GFP+^ cells among Tfh cells. (B) Frequency of IL-4^GFP+^ cells among Tfh cells. (C) Flow cytometry analysis to identify GATA3^+^ (Top), ROR-γt^+^ (Middle), T-bet^+^ (Bottom) cells among Tfh cells. (D) Frequency of GATA3^+^ (Top), ROR-γt^+^ (Middle), T-bet^+^ (Bottom) cells among Tfh cells. (E) gMFI of FITC 2-NBDG on Tfh cells after *ex vivo* incubation with FITC 2-NBDG. (F) gMFI of mitotracker green of Tfh cells. (G) gMFI of mitotracker red of Tfh cells. Data are representative of three independent experiment, ns=not significant, ****p<0.001, *p<0.05, unpaired t test.

**Supplementary Figure 3 (related to figure 3).**
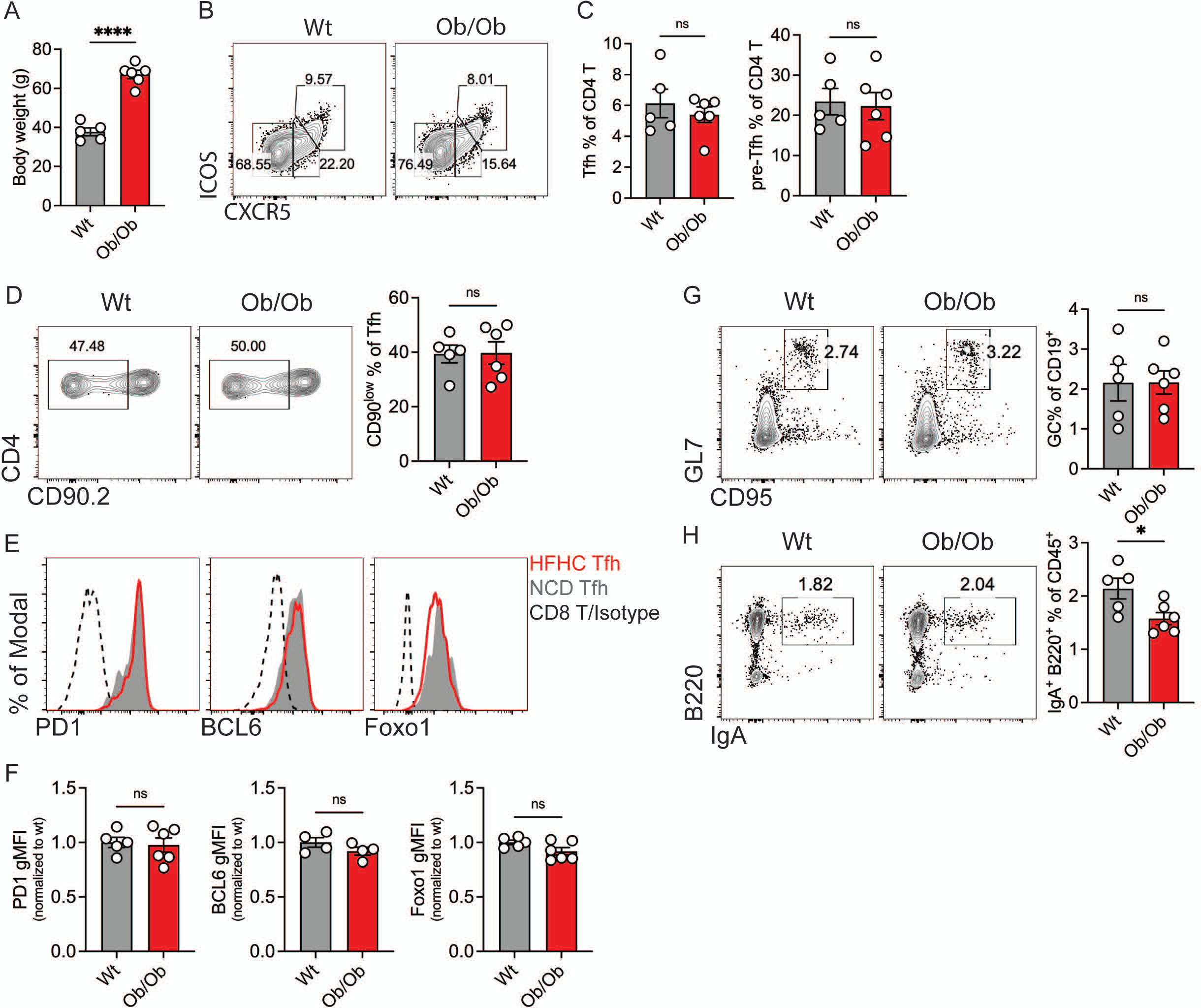
(A) Body weight Wt and Ob/Ob mice. (B) Representative flow cytometry analysis to identify Tfh (CXCR5^high^ ICOS^high^) and Pre-Tfh (CXCR5^intermediate^ ICOS^intermediate^) cells among Foxp3^−^ CD4 T cells. (C) Frequency of Tfh (Left) and Pre-Tfh (Right) cells among total CD4 T cells. (D) Representative flow cytometry analysis to identify CD90^low^ cells among Tfh cells (Left), frequency of CD90^low^ cells among Tfh cells (Right). (E) Representative flow cytometry histogram of PD1 (Left), BCL6 (Middle), FOXO1 (Right) expression in HFHC Tfh cells, NCD Tfh cells. (F) gMFI of PD1 (Left), BCL6 (Middle), FOXO1 (Right) in HFHC Tfh cells, NCD Tfh cells. (G) Representative flow cytometry analysis to identify GC (CD95^+^ GL7^+^) among CD19^+^ B cells (Left), frequency of GC among CD19^+^ B cells (Right). (H) Representative flow cytometry analysis to identify IgA^+^ B cells (IgA^+^ B220^+^) among total live CD45^+^ cells (Left), frequency of IgA^+^ B among total live CD45^+^ cells (Right). Data are representative of three independent experiment, ns=not significant, ****p<0.001, unpaired t test.

**Supplementary Figure 4 (related to figure 3).**
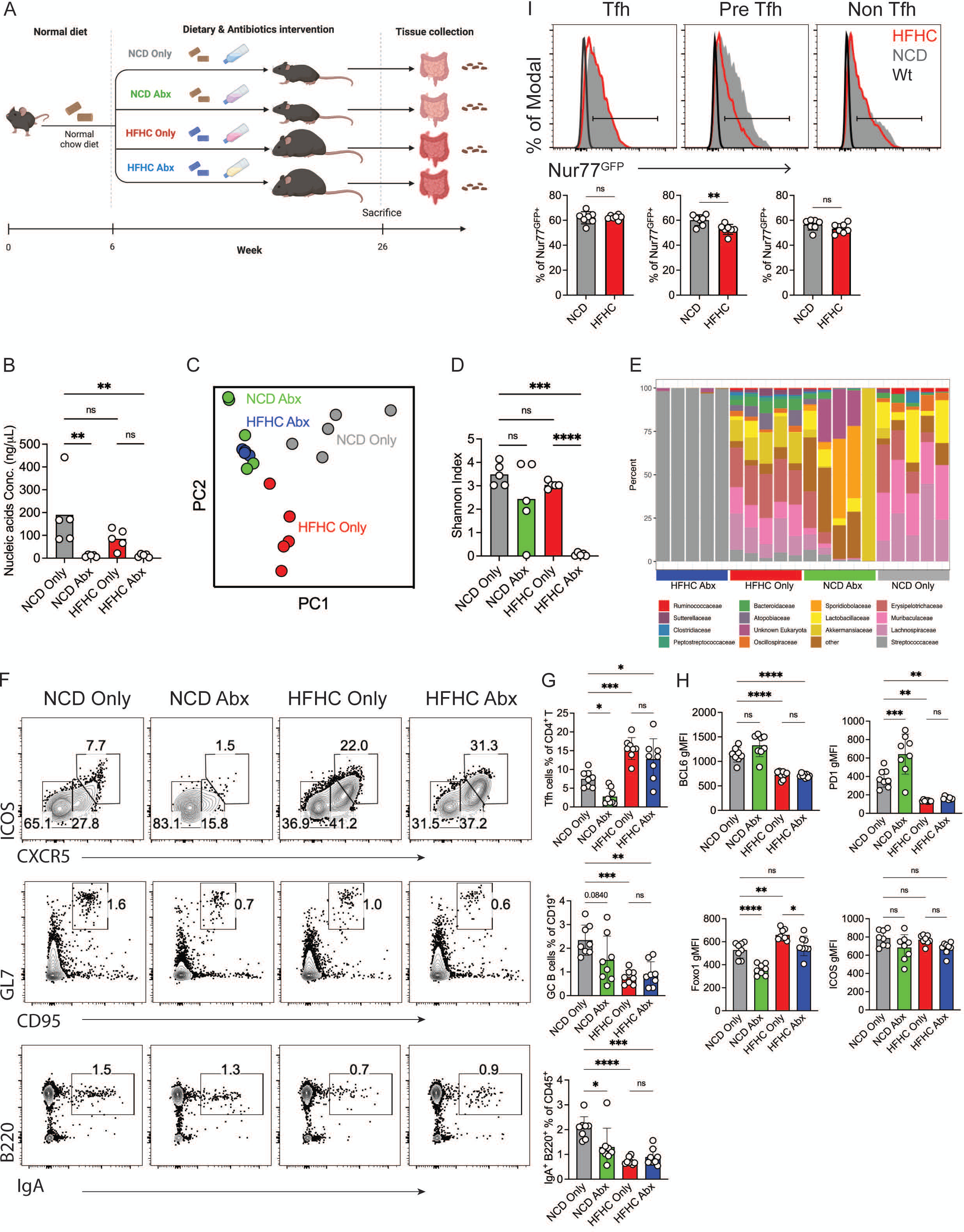
(A) Experimental scheme. (B) Concentration of nucleic acids extracted from fecal pellets of NCD or HFHC mice treated with or without antibiotics. (C) PCA plot of 16S rRNA sequencing data of intestinal microbiota from NCD or HFHC mice treated with or without antibiotics. (D) Shannon diversity index of 16S rRNA sequencing data of intestinal microbiota from NCD or HFHC mice treated with or without antibiotics. (E) Composition of microbiota community from NCD or HFHC mice treated with or without antibiotics. (F) Representative flow cytometry analysis to identify Tfh cells (Top), GCs (Middle) and IgA^+^ B cells (Bottom) among Foxp3^−^ CD4 T cells, CD19^+^ B cells and total live CD45^+^ cells, respectively. (G) Frequency of Tfh cells (Top), GCs (Middle) and IgA^+^ B cells (Bottom) among Foxp3^−^ CD4 T cells, CD19^+^ B cells and total live CD45^+^ cells, respectively. (H) gMFI of BCL6 (Top left), PD1 (Top right), FOXO1 (Bottom left), ICOS (Bottom right) expression in Tfh cells. (I) Representative flow cytometry histogram to identify Nu77^GFP+^ cells among Tfh (Top left), Pre-Tfh (Top middle) and Non-Tfh (Top right) cells in HFHC and NCD mice, frequency of Nu77^GFP+^ cells among Tfh (Bottom left), Pre-Tfh (Bottom middle) and Non-Tfh (Bottom right) cells in HFHC and NCD mice. Data are representative of three independent experiment, ns=not significant, ****p<0.001, ***p<0.005, **p<0.01, *p<0.05, unpaired t test or ANOVA.

**Supplementary Figure 5 (related to figure 4).**
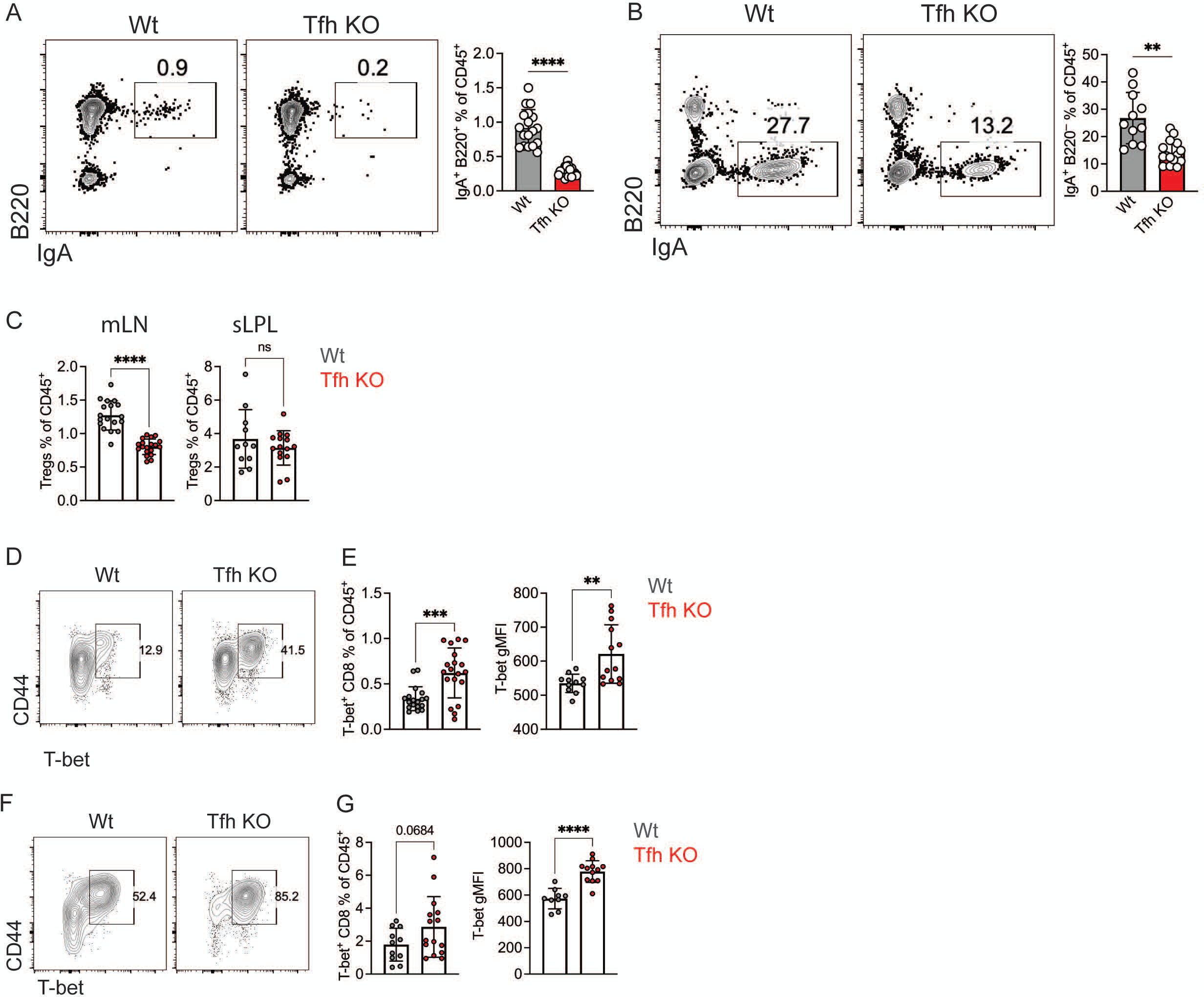
(A) Representative flow cytometry analysis to identify IgA^+^ B cells (IgA^+^ B220^+^) among total live CD45^+^ cells in PPs (Left), frequency of IgA^+^ B cells (IgA^+^ B220^+^) among total live CD45^+^ cells in PPs (Right). (B) Representative flow cytometry analysis to identify IgA^+^ B cells (IgA^+^ B220^−^) among total live CD45^+^ cells in small intestine LPL (Left), frequency of IgA^+^ B cells (IgA^+^ B220^−^) among total live CD45^+^ cells in small intestine LPL (Right). (C) Frequency of Tregs among total live CD45^+^ cells in mLN (Left) and small intestine LPL (Right). (D) Representative flow cytometry analysis to identify pro-inflammatory CD8 T cells (T-bet^+^ CD44^+^) in PPs. (E) Frequency of pro-inflammatory CD8 T cells (T-bet^+^ CD44^+^) among total live CD45^+^ cells in PPs (Left), gMFI of T-bet in pro-inflammatory CD8 T cells (T-bet^+^ CD44^+^) in PPs (Right). (F) Representative flow cytometry analysis to identify pro-inflammatory CD8 T cells (T-bet^+^ CD44^+^) in small intestine LPL. (G) Frequency of pro-inflammatory CD8 T cells (T-bet^+^ CD44^+^) among total live CD45^+^ cells in small intestine LPL (Left), gMFI of T-bet in pro-inflammatory CD8 T cells (T-bet^+^ CD44^+^) in small intestine LPL (Right). Data are representative of three independent experiment, ns=not significant, ****p<0.001, ***p<0.005, **p<0.01, unpaired t test.

**Supplementary Figure 6 (related to figure 5).**
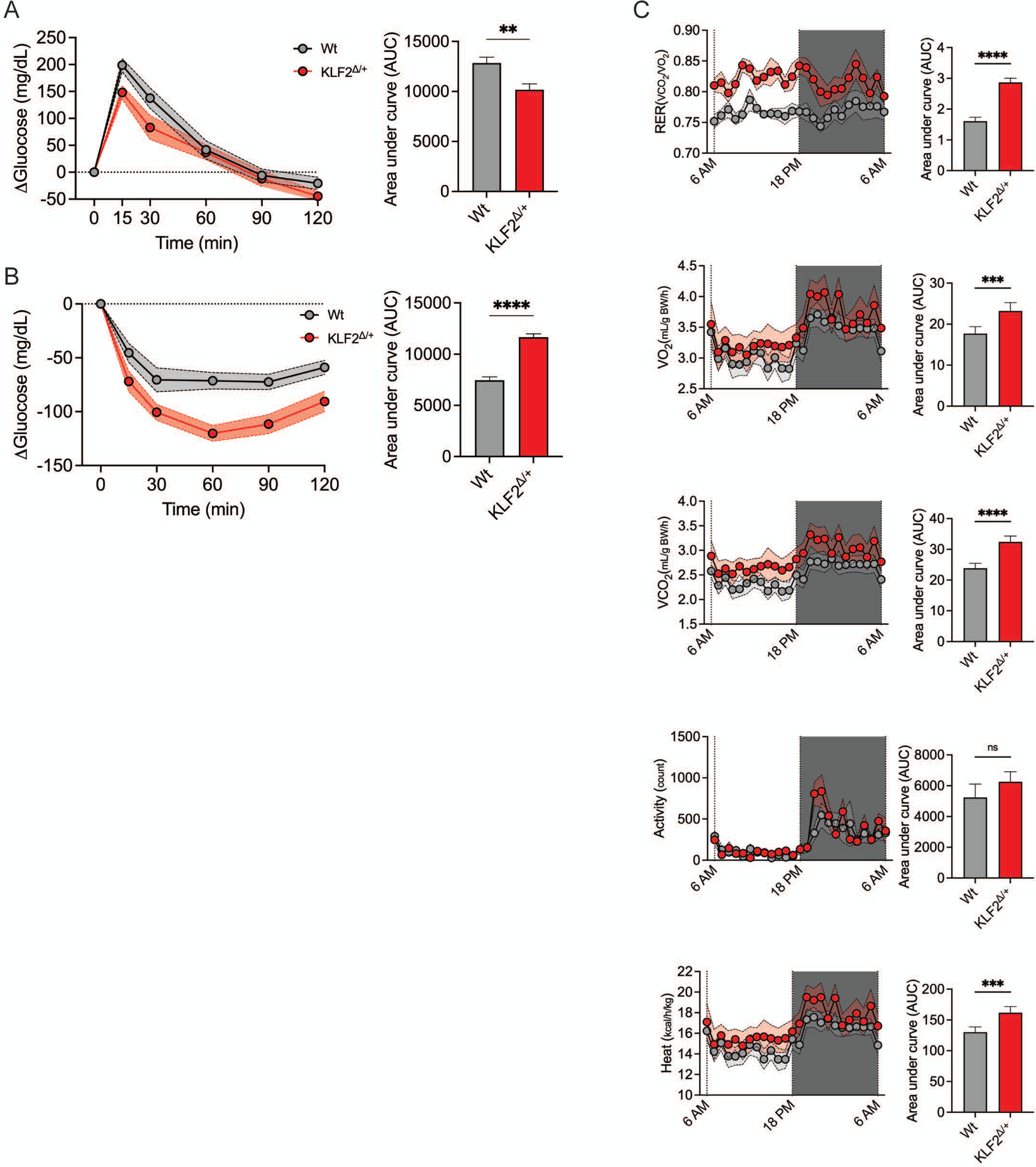
(A) Change of glucose level in blood during glucose tolerance test (GTT) (Left), corresponding area under curve (AUC) calculated from GTT assay (Right). (B) Change of glucose level in blood during insulin tolerance test (ITT) (Left), corresponding area under curve (AUC) calculated from ITT assay (Right). (C) Hourly respiratory exchange ratio (RER) for 24 hours (1^st^ column left), and corresponding AUC (1^st^ column right); Hourly oxygen consumption for 24 hours (2^nd^ column left), and corresponding AUC (2^nd^ column right); Hourly carbon dioxide production for 24 hours (3^rd^ column left), and corresponding AUC (3^rd^ column right); Hourly ambulatory activity for 24 hours (4^th^ column left), and corresponding AUC (4^th^ column right); Hourly energy expenditure for 24 hours (5^th^ column left), and corresponding AUC (5^th^ column right).

